# Sensorimotor adaptation of vocal pitch is severely impaired in cerebellar ataxia

**DOI:** 10.1101/2025.10.10.681608

**Authors:** Anneke Slis, Benjamin Parrell

## Abstract

Sensory errors, mismatches between predicted sensory outcomes of movement and reafferent sensory feedback, drive changes in the feedforward control of future motor behavior that correct for those errors. Across a wide variety of motor behaviors, individuals with cerebellar damage show impairments in these corrections, strongly suggesting a key role of the cerebellum in sensorimotor adaptation. However, the extent to which the cerebellum is involved in controlling vocal pitch is currently unknown. Crucially, vocal pitch differs in several ways from other systems that suggest it relies more on feedback than feedforward control. Adaptation itself also differs in vocal pitch: rather than the gradual build-up/decay of learning seen in other systems, pitch adaptation and de-adaptation are almost immediate. Together, this questions whether adaptation in vocal pitch relies on the same mechanism as other motor domains. Here, we test the hypothesis that the cerebellum underlies sensorimotor adaptation in vocal pitch, testing the domain-generality of this neurocomputational process. In both sustained vocalization and a more natural word production task, individuals with cerebellar ataxia fail to adapt to external auditory perturbation of vocal pitch. The complete lack of adaptation observed, compared to the impaired but present adaptation seen in other systems, suggest that the cerebellum plays an especially critical role in maintaining accurate control of vocal pitch. Conversely, we failed to detect a previously observed increase in online compensation to vocal pitch errors in ataxia, potentially suggesting this may be an idiosyncratic change in control rather than a common trait in this population.

**Significance statement:** When exposed to external perturbations of vocal pitch, individuals with cerebellar ataxia fail to adapt their produced pitch to oppose the perturbation. These results highlight the critical role of the cerebellum in driving adaptation across motor domains, even for motor behaviors that are thought to rely more on feedback compared to feedforward control, such as pitch. Conversely, and despite a large cohort, we failed to replicate previous findings of enhanced online corrections for pitch perturbations in ataxia, suggesting that any such increases are likely due to individual-specific changes in motor function in response to cerebellar-induced control deficits rather than being directly related to cerebellar damage itself.

## Introduction

Vocal pitch is an important component of spoken communication, conveying pragmatic and, in some languages, lexical information. Auditory feedback plays a critical role in accurately producing pitch, shown experimentally when speakers adjust their vocalisations to counter external auditory perturbations of their perceived pitch (for a review, see Parrell & Houde, 2019). These adjustments occur over multiple timescales: speakers reactively fine-tune their pitch during ongoing vocalizations (Burnett et al., 1998; Hain et al., 2000) and also adjust their pitch in an anticipatory fashion on future trials (Alemi et al., 2020; Behroozmand & Sangtian, 2018; Jones & Munhall, 2002, 2002, 2000, 2005).

This latter process, sensorimotor adaptation, is thought to be driven by comparing sensory predictions to incoming sensory feedback; any discrepancies lead to updates to the internal models used in feedforward control (Parrell & Houde, 2019; Shadmehr & Krakauer, 2008). Sensorimotor adaptation has been observed across multiple motor domains and is thought to be driven largely by the cerebellum, reflected by impaired adaptation across multiple motor domains in individuals with cerebellar ataxia (CA), including oculomotor control, reaching, locomotion, and speech (Flament et al., 1996; Manto et al., 2012; Martin et al., 1996; Morton & Bastian, 2006; Parrell et al., 2021; Smith & Shadmehr, 2005; Tseng et al., 2007). Conversely, individuals with CA exhibit intact feedback mechanisms in reaching, walking, and oral articulation, differing little from neurobiologically healthy individuals (NH) in the magnitude of online compensatory responses (Morton & Bastian, 2006; Parrell et al., 2021; Zimmet et al., 2020).

When exposed to pitch perturbations, however, individuals with CA show larger compensatory responses than NH speakers (Houde et al., 2019; Li et al., 2019). While the exact mechanism underlying these enlarged responses in CA is not clear, it suggests that pitch control may differ in fundamental ways from both oral articulation and limb control. Indeed, relative to other motor behaviors, controlling pitch is hypothesized to rely more directly on sensory feedback, as shown, for example, by the rapid decline of vocal control in postlingually deaf individuals compared to the much slower changes in oral articulation in the same population (Cowie & Douglas-Cowie, 1992; Lane & Webster, 1991; Perkell et al., 1997). Relatedly, producing a pitch contour – that is, realizing a sequence of high and low pitches to indicate prosodic events – relies on achieving pitch values relative to previous pitch targets, rather than on absolute values as in the case of spatial targets (see e.g. Wagner & Watson, 2010). Most strikingly, sensorimotor adaptation to pitch perturbations also differs in several important ways from patterns observed in other motor domains. Adaptation in reaching, walking, eye movements, and oral articulation shows a gradual build-up over multiple trials, a gradual return to baseline behavior once the perturbation is removed, and is consistently observed across studies. Conversely, pitch adaptation is sudden, stabilizes after only a few trials, disappears rapidly at perturbation removal (Alemi et al., 2020; Behroozmand & Sangtian, 2018; Jones & Keough, 2008; Keough et al., 2013), and varies substantially in magnitude and presence across and within studies (Kapsner-Smith et al., 2024).

These differences in pitch control mechanisms relative to other motor behaviors raise questions about whether sensorimotor adaptation to pitch perturbations relies on a similar cerebellar learning circuit as in other motor domains. To answer this question, we assess whether adaptation to perturbations of vocal pitch is impaired in individuals with CA. Additionally, we measure online compensatory responses to pitch perturbations in the same cohort to 1) replicate previous findings of enhanced compensatory responses in CA compared to NH and 2) examine the potential relationship between impairments in sensorimotor adaptation and increases in feedback-based compensatory responses in CA, based on the hypothesis that enhanced compensatory responses indicate reorganization in control subsequent to feedforward impairments (Houde et al., 2019). To fully assess feedback control of pitch, we assess compensation to perturbations delivered at vowel midpoint (typical of previous studies) as well as vowel onset.

## Materials and Methods

### Participants

43 individuals with CA (17 M, 26 F; mean age = 51, *SD* = 14, age range: 21-74) and 48 age-matched NH individuals (18 M, 30 F; mean age = 53, *SD* = 16, age range: 22-80) participated in the study. All participants completed Experiments 1 and 2. A subset of these participants also completed Experiment 3 (CA: 30 total; 14 M, 16 F; mean age = 50, *SD* = 15, age range: 21-75; NH: 27 total; 11 M, 14 F; mean age = 50, *SD* = 16, age range: 22-77) and Experiment 4 (CA: 17 total; 5 M; 12 F; mean age =51, *SD* =14, age range: 21-73; NH: 17 total; 9 M, 8 F; mean age = 42, *SD* = 15, age range: 22-77. All participants were native American English speakers, and reported no hearing, speech, language, or neurological impairments other than cerebellar ataxia. The etiology of individuals with CA was heterogeneous, but the majority received a diagnosis of some type of spinocerebellar ataxia from their neurologist (Table I). A subset of these individuals had also received a confirmation of spinocerebellar ataxia type by genetic testing. All participants were assessed on their performance in executive functioning, spatial cognition, language, and affect, using the Cerebellar Cognitive Affective Syndrome scale (CCAS). In addition, cognitive functioning was evaluated using the Montreal Cognitive Assessment (MoCA). Individuals with CA received additional assessments to score the severity of their ataxia with the Scale for Assessment and Rating of Ataxia (SARA) and to evaluate whether neurologic issues unrelated to ataxia were present with the Inventory of Non-Ataxia Signs (INAS). Participant hearing was tested with a Hughston Westlake hearing screening but, given the prevalence of high-frequency hearing loss in older adults, participants were not excluded based on their hearing thresholds. No participant used assisted hearing devices. Participants were monetarily compensated for their participation. All procedures were approved by an Institutional Review Board at the University of California, San Francisco.

**Table I:** Participant details for individuals with CA. SARA: higher scores indicate more severe ataxia (maximum score 40); CCAS raw: lower scores indicate more severe effects of ataxia (maximum score 120); CCAS Pass/Fail: number of failed tests: 0 normal, 1 Possible CCAS, 2 Probable CCAS, 3 or more Definite CCAS (maximum 10). INAS: non-ataxia signs, higher scores, more severe (maximum 16). Two participants were excluded (indicated in gray): CA024 was not able to sustain pitch for the required time and CA118 did not complete the full set of experiments.

| participant | age | gender | subtype | MoCA | SARA | CCAS raw | CCAS # pass/fail | INAS |
| --- | --- | --- | --- | --- | --- | --- | --- | --- |
| **Experiments 1 and 2** |  |  |  |  |  |  |  |  |
| CA001 | 46 | F | AOA2 | 26 | 24 | 95 | 2 | 6 |
| CA031 | 65 | F | SCA6 | 28 | 16 | 109 | 0 | 6 |
| CA073 | 74 | M | SCA6 | 23 | 9 | 91 | 2 | 3 |
| CA086 | 70 | F | CANVAS | 23 | 20.5 | 88 | 4 | 8 |
| CA085 | 52 | F | tumor | 25 | 9 | 92 | 0 | 5 |
| CA091 | 54 | F | SCA14 | 29 | 8.5 | 103 | 0 | 2 |
| CA092 | 39 | F | SCA3 | 29 | 17.5 | 97 | 0 | 9 |
| CA094 | 52 | F | SCA5 | 28 | 13 | 90 | 2 | 5 |
| CA062 | 32 | M | SCA2 | 23 | 23.5 | 77 | 5 | 9 |
| CA060 | 69 | M | SCA2 | 23 | 10 | 83 | 3 | 2 |
| CA095 | 44 | F | SCA2 | 28 | 7.5 | 90 | 2 | 7 |
| CA024 | 55 | F | SCA3 | 25 | 18.5 | 81 | 3 | 8 |
| CA089 | 60 | F | SCA8 | 28 | 13.5 | 98 | 0 | 3 |
| **Experiments 1, 2, and 3** |  |  |  |  |  |  |  |  |
| CA088 | 25 | F | SCA | 25 | 11 | 77 | 3 | 2 |
| CA103 | 63 | M | SCA8 | 26 | 15.5 | 92 | 1 | 5 |
| CA093 | 39 | M | SCA8 | 26 | 2.5 | 99 | 1 | 3 |
| CA098 | 59 | M | SCA6 | 25 | 10 | 106 | 1 | 3 |
| CA106 | 24 | M | SCA16 | 27 | 20.5 | 81 | 4 | 1 |
| CA096 | 62 | M | SCA3 | 21 | 13 | 75 | 5 | 10 |
| CA115 | 34 | M | SCA2 | 30 | 11.5 | 111 | 0 | 7 |
| CA109 | 52 | M | idiopathic | 24 | 27 | 74 | 3 | 6 |
| CA107 | 64 | M | SCA2 | 27 | 11 | 91 | 1 | 5 |
| CA111 | 41 | F | SCA3 | 27 | 9 | 90 | 1 | 2 |
| CA122 | 73 | F | SCA8 | 26 | 19 | 79 | 4 | 7 |
| CA065 | 44 | M | SCA2 | 28 | 6.5 | 102 | 1 | 1 |
| CA116 | 75 | F | SCA3 | 26 | 1 | 102 | 2 | 3 |
| CA069 | 39 | F | SCA1 | 29 | 1 | 109 | 0 | 4 |
| **Experiments 1, 2, 3 and 4** |  |  |  |  |  |  |  |  |
| CA105 | 65 | F | SCA27 | 27 | 1 | 107 | 0 | 1 |
| CA110 | 60 | M | SCA2 | 25 | 12 | 95 | 2 | 5 |
| CA112 | 32 | F | SCA5 | 29 | 2 | 111 | 0 | 2 |
| CA114 | 52 | F | SCA4 | 27 | 1 | 100 | 2 | 0 |
| CA121 | 35 | M | SCA4 | 23 | 10.5 | 89 | 1 | 3 |
| CA119 | 35 | F | SCA3 | 30 | 1 | 114 | 0 | 1 |
| CA124 | 68 | F | SCA2 | 24 | 17.5 | 84 | 1 | 5 |
| CA043 | 52 | F | SCA2 | 27 | 10 | 105 | 1 | 13 |
| CA008 | 71 | F | SCA3 | 27 | 30 | 93 | 1 | 21 |
| CA126 | 43 | M | SCA8 | 22 | 15 | 65 | 6 | 13 |
| CA125 | 53 | M | SCA8 | 25 | 9.5 | 90 | 1 | 11 |
| CA030 | 63 | F | SCA3 | 28 | 6 | 108 | 0 | 2 |
| CA108 | 47 | F | SCA2 | 30 | 14 | 115 | 0 | 5 |
| CA118 | 43 | M | SCA3 | 21 | 22 | 62 | 7 | 10 |
| CA123 | 56 | F | SCA5 | 28 | 12.5 | 92 | 2 | 2 |
| CA127 | 21 | F | SCA13 | 25 | 12 | 94 | 2 | 6 |

### Software/ hardware

Participants’ speech was recorded at 48 Khz using a Sennheiser MKE 600 shotgun microphone and a Focusrite scarlett 2i2 sound card, and downsampled to 16 kHz and processed (and on some trials perturbed), using Audapter (Cai et al., 2008, 2014; Tourville et al., 2008). The resulting output was played back to participants at 80 dB SPL in near real-time over closed-back, circumaural Beyerdynamic DT 770 PRO 250-Ohm headphones. The output signal was calibrated each time before the start of the experiment with a CM-140 Checkmate SPL Sound Pressure Meter, confirming that masking noise, played back via the headphones, had an intensity of ∼60 dBA. While the speech signal was calibrated before the start of the experiment to play at ∼80dB, this calibration set a constant feedback gain applied to the input signal, such that if the speaker changed their intensity during the experiment, the intensity of the feedback signal changed accordingly. Thus, to maintain the desired output level, the output intensity level was monitored during the study and, in cases where a participant consistently spoke either too loud or too soft (for example in case of fatigue), the microphone level was adjusted to maintain the ∼80 dB output in the signal that was fed back to the headphones.

All pitch shifts were implemented using the time-domain shifting algorithm in Audapter with a buffer length of 64 samples, which left the remainder of the spectral envelope, including vowel formants, unaffected. The pitch shift algorithm resulted in a measured delay of ∼38 ms in the auditory feedback signal delivered to participants, which has been shown not to affect the magnitude of vocal responses (Weerathunge et al., 2020). To determine an estimate of the a-priori frequency range to accurately track pitch in real time, the average fundamental frequency of vocal fold vibration (*f0*, the physical signal underlying the auditory percept of pitch) was estimated in a brief calibration period before each experiment, using the “pitch” function in MATLAB (The Mathworks, Inc, 2022). Boundaries were generally set 20% above and 20% below the average *f0*; some speakers required a larger range (2 NH/11 CA: 25%, 5 NH/4 CA: 30%, and 1 CA: 40%) due to high variability in their produced pitch. Subsequently, during each study, these upper and lower *f0* values were updated after each trial using the same percentage window around the average *f0* of that specific trial. This was done to account for any drift in *f0* over time, which has been observed previously in studies of adaptation and which is also sometimes observed when producing a series of repeated vocalizations (Alemi et al., 2021; Jones & Munhall, 2000).

### Experimental design

Data were collected as part of a larger battery of experiments lasting approximately 9 hours in total, conducted over either 2 days (for individuals with CA) or 1-3 days (for NH individuals). All participants first completed Experiment 1 (examining pitch adaptation), followed by Experiment 2 (examining online, reflexive compensation to sustained mid-utterance perturbations of vocal pitch). Based on the adaptation and compensation responses from the first 13 CA and 21 NH participants, a subset of participants (30 CA and 27 NH) additionally completed Experiment 3, examining reflexive compensation to transient perturbations of vocal pitch applied at vocalization onset. Finally, a smaller subset (17 CA and 17 NH) participated in Experiment 4, examining reflexive compensation to transient mid-utterance pitch perturbations. Data from one NH participant (Exps 1-4) were disregarded, due to technical difficulties with the software during collection of the unperturbed production of the sustained vowel, leaving a total of 47 NH participants. Data from two individuals with CA (one participating in Exps 1-2; the other in Exps 1-4) were disregarded because these speakers were unable to produce extended phonation of sufficient duration, resulting in a final data set containing 41 CA individuals.

The order of the experiments was identical for all participants. Experiment 1, examining adaptation, preceded the 3 compensation experiments (Experiment 2, 3, and 4). Experiment 1 was administered on a separate day than Experiment 2, 3, and 4 for most participants, except those NH individuals who chose to complete all experiments in a single day (14 NH individuals) and a few individuals with CA who, due to time constraints, did not complete the adaptation study on the first day of their visit (13 CA individuals). Experiments 3 and 4 always followed Experiment 2 (in that order). Although Experiments 2, 3 and 4 were completed on the same day, they were always separated by other experiments in the larger battery unrelated to vocal pitch control.

During each experiment, the participants were comfortably seated in front of a computer screen. They were cued to produce speech by a written stimulus appearing on the screen. For all except two experiments (as noted below for part of Experiment 1), participants produced a sustained vowel (/ɑ/ as in “hot”), indicated with the written stimulus “ah”. Participants were instructed to produce this vowel as soon as they saw the stimulus appear on the screen and to vocalize until the stimulus disappeared (after 1.4s), keeping their pitch as stable as possible. Pilot testing showed that this duration resulted in vocal productions of ∼1 s. The amplitude, onset time, and duration of vocal productions were monitored on every trial. On trials with vocalization durations < 1s, participants were reminded to produce the vowel until it disappeared from the screen. When participants began vocalizing > 400 ms after stimulus onset, they were reminded to start sooner. On trials where the speech amplitude was below a preset threshold, they were prompted to speak a little bit louder. The inter-trial duration was 1.25 seconds with 0-0.25 seconds of additional random jitter.

#### Experiment 1

Experiment 1 (Figure 1B) tested adaptive responses to consistent auditory pitch perturbations both during sustained vocalization and, in a separate session, during word production. To account for the fact that pitch has been shown, in some cases, to drift over time, especially during sustained vocalization (Alemi et al., 2021; Jones & Munhall, 2000), we included an additional unperturbed control session for both sustained vocalization and spoken word stimuli. This resulted in four total sessions in this study. During the first two sessions, participants produced the word “bod” (/bɑd/), cued by the stimulus “bod” on the screen. During the second two sessions, participants produced a sustained vowel /ɑ/, cued by the stimulus “ah”. Each session consisted of 80 trials.

**Figure 1.**
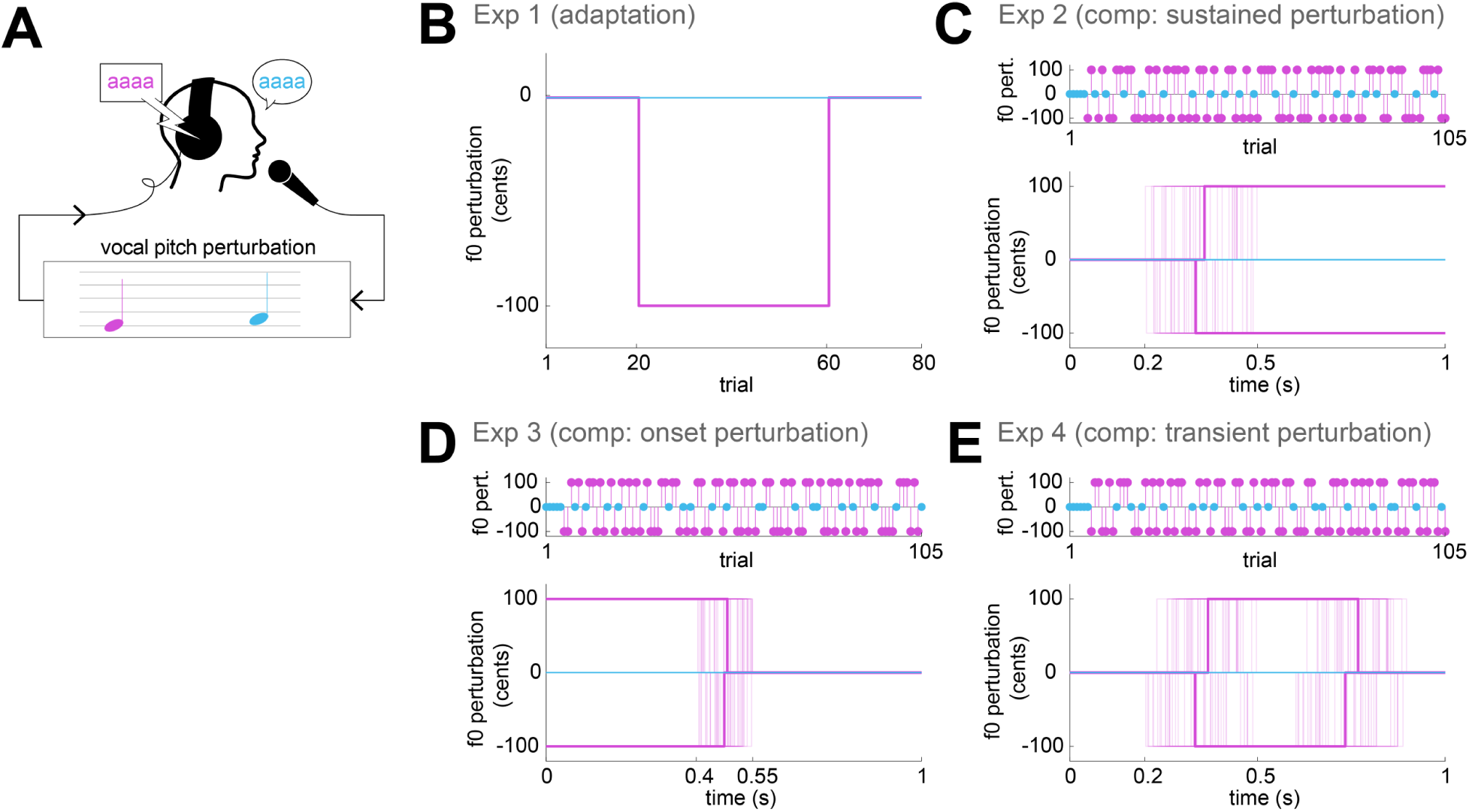
Schematic of experimental design. In all cases, unperturbed productions are shown in cyan and productions with a vocal pitch perturbation in magenta. **A**: Illustration of pitch perturbation. The pitch of participants’ produced vocalizations were shifted down by 100 cents and played back to them over headphones in essentially real time (∼38 ms delay). **B**: Perturbation schedule in Experiment 1, examining across-trial sensorimotor adaptation. The perturbation session (magenta) consisted of a 20-trial baseline phase with no pitch perturbation, a 40-trial exposure phase with a -100 cent perturbation, and a 20-trial washout phase with no perturbation. The control session (cyan) had veridical feedback throughout. **C:** Design of Experiment 2, examining within-trial compensatory responses to sustained mid-vocalization perturbations. The perturbation direction was randomized across trials (top). Within each trial, perturbations started between 200 and 500 ms after vocalization onset and remained on until the end of the trial. Thick magenta lines show the mean time course of the perturbation; thin lines show individual trials. Example shown from an individual participant. **D:** Design of Experiment 3, examining compensatory responses to transient perturbations beginning at the onset of vocalization, and removed between 400 and 550 ms after vocalisation onset. Example perturbation data as in (C) **E**: Design of Experiment 4, examining compensatory responses to transient 400-ms perturbations beginning 200-500 ms after vowel onset. Example perturbation data as in (C).

Sessions 1 and 3 (“perturbed” sessions) consisted of a 20-trial baseline phase with no perturbation, a 40-trial exposure phase where pitch was shifted down by 100 cents and a 20-trial washout phase with veridical feedback. A shift of 100 cents was chosen because larger shifts have been shown to evoke more following responses, potentially because the shift is treated as an external target rather than as a sensory prediction error (Burnett et al., 1998). Sessions 2 and 4 (“unperturbed” sessions) contained the same number of trials as the perturbed sessions (80), with no pitch perturbation applied at any point. Prior to the first and third session, participants produced 5 practice trials to make sure they understood and produced the target stimuli correctly. These practice trials were always unperturbed. A brief self-timed break occurred every 14 trials. The total duration of Experiment 1, including all four sessions, was ∼20 minutes.

#### Experiment 2

Experiment 2 (Figure 1C) consisted of 105 trials during which the participants produced a sustained vowel /ɑ/. The first 5 trials received no perturbation. During 80 of the remaining 100 trials, pitch was shifted by ±100 cents (40 trials each direction) starting 200-500 ms after vocalisation onset (randomly applied across trials) and continuing throughout the trial (see figure 1B). On the remaining 20 trials, the participant received veridical auditory feedback. The 100 trials containing the 3 conditions (40 shift up, 40 shift down, 20 unperturbed) were randomized across trials. A short self-timed break occurred every 21 trials. The total study took ∼10 minutes to complete.

#### Experiments 3 and 4

The format of Experiment 3 (Figure 1D) and Experiment 4 (Figure 1E) were essentially identical to Experiment 2, except for the timing of the pitch perturbation. While the perturbation in Experiment 2 randomly started 200-500 ms after the onset of vocalization and was applied through the remainder of the trial, the perturbation in Experiment 3 always began at the onset of vocalisation and was randomly removed after 400-550 ms (drawn from a normal distribution across trials). In Experiment 4, the perturbation began 200-500 ms after vocalization onset (drawn from a normal distribution across trials), as in Experiment 2, but was removed after 400 ms.

### Data analysis

We extracted *f0* using pitch tracking algorithms from PRAAT (Boersma & Weenink, 2020) embedded within Wave_Viewer (Niziolek & Houde, 2015), a custom-built speech analysis package in MATLAB (The Mathworks, Inc, 2022). The pitch floor in PRAAT was set at a default value of 75 Hz and the upper threshold was set at 200 Hz for lower-pitched voices and 300 Hz for higher-pitched voices. The default voicing threshold was 0.45. When these values resulted in errors in pitch tracking (for example resulting in large jumps from sample to sample or pitch halving), the PRAAT pitch tracking parameters were changed on a trial-by-trial basis to remove these errors. Pitch floors were adjusted when pitch halving was observed. As different pitch floors result in different sampling rates in PRAAT, all trials were resampled to a uniform sampling rate of 100 samples per second. Trials with irresolvable tracking errors in the target window used for analysis were excluded. Trials on which participants began phonation prior to the onset of the recording (equivalent to beginning prior to the stimulus presentation) were also excluded. For Experiments 2-4, trials with a duration less than 1000 ms were excluded, as this was the minimum duration necessary to capture the full dynamics of compensation (pitch perturbation began as late as 500 ms after vowel onset, with 500 ms needed to measure a response plateau). The total number of excluded trials was: Experiment 1: 2.54 %, *SD* = 4.54 %; range 0-26 % across participants; Experiment 2: 1.83 %, *SD* = 5.04 %; range 0-32%; Experiment 3: 0.80 %, *SD* = 1.44 %, range 0-7.76 %; Experiment 4: 0.63 % *SD* = 1.22 %, range 0-4.74%.

#### Experiment 1

To isolate adaptive changes to pitch perturbations, the *f0* for each trial was extracted from a window 50-120 ms after vowel onset to avoid 1) irregularities in pitch production and pitch tracking that frequently occur during the first few cycles of vocal fold vibration (Heller Murray & Stepp, 2020; Kapsner-Smith et al., 2024) and 2) potential online compensatory mechanisms, which typically start ∼150 ms after the onset of a pitch perturbation (Burnett et al., 1998). Pitch values for all trials were transformed to a cents scale to be able to compare responses across participants with different *f0* values (cents are a logarithmic scale used to measure pitch intervals). First, the median *f0* of each of the 10 final trials during the baseline phase for each session was calculated over the 50-120 ms window, giving one *f0* value for each trial. Next, for each session, a baseline value was calculated as the median of these trials. This baseline value was then used to convert the pitch trajectories of all the 80 trials in that session to a cent scale using the formula 1200*log_2_(*f0*/*f0*_baseline_). Next, a single pitch value for each trial was calculated as the median pitch, expressed in cents, in the target time window of 50-120 ms. To account for the observable drift in pitch observed in the unperturbed sessions, especially during sustained vowel production, *f0* values of all trials in the perturbation sessions were corrected by subtracting the moving average across 15 trials produced during the corresponding unperturbed session (Figure 2A & 2C).

**Figure 2:**
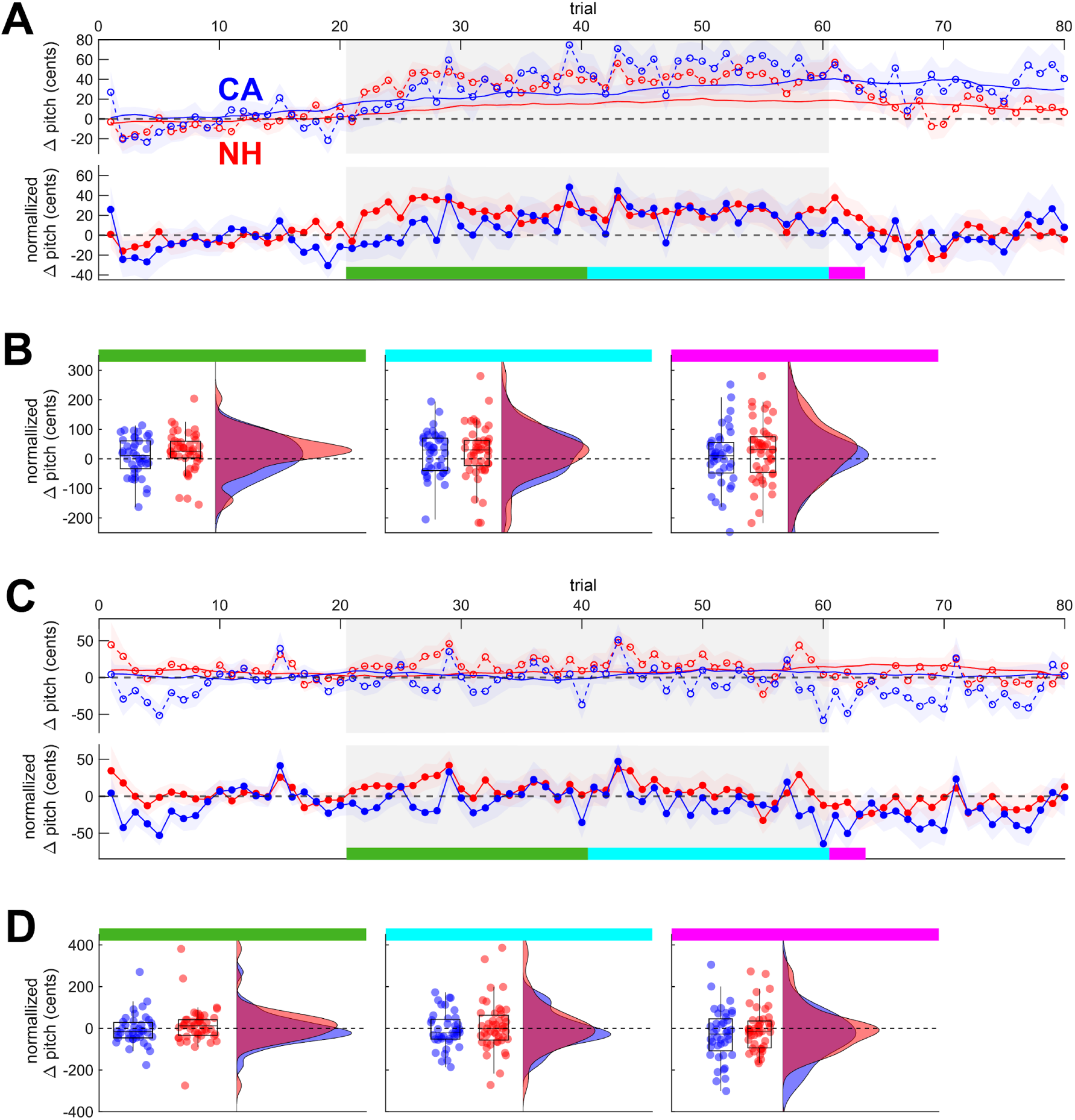
Adaptation results from Experiment 1. In all panels, blue indicates values for CA speakers and red indicates values for NH speakers; **A**: Top panel: Pitch values during vowel production in the unperturbed (solid lines) and perturbed (dashed lines) sessions. Bottom Panel: pitch values in the perturbed session, normalized by a moving average of the unperturbed session, during vowel production. Shaded areas represent standard error. Gray shading indicates exposure phase. Dark green, cyan, and magenta shading represent trials used to analyze the early exposure, late exposure, and washout phases, respectively. **C**: Normalized pitch values produced in the perturbed session of the vowel production study in the early exposure phase (left), late exposure phase (middle), and washout phase (right). . Dots and density plots represent participant means.. **C-D**: As for A-B, showing data for word production.

For word production, pitch trajectories in English show a large decline in pitch over the course of the vowel during a trial. It may be that only a part of this pitch contour is adapted (for example, the first high pitch target) rather than the full trajectory. To assess this possibility, we conducted an additional analysis using Generalized Additive Mixed Models (GAMMs) to examine the full time course of pitch values (10-250 ms after vowel onset), comparing the final 10 trials in the baseline phase with early exposure (first 10 trials), late exposure (final 10 trials) and washout phases (first 3 trials) across CA and NH groups. Given that minimal changes in pitch were observed over the course of the unperturbed session (Figure 2C), pitch trajectories for this analysis were not normalized to the unperturbed session. Individual models were constructed to test for adaptation within each experimental group separately and, for each phase other than baseline, to test for differences between the CA and NH groups. For the between-group models, trajectories in each phase were first normalized by subtracting the mean baseline trajectory, on a participant specific basis, to conservatively account for any potential differences in baseline trajectories between groups (although visual inspection of the group-means shows broadly similar baseline behavior). The input to these models thus reflect the change in pitch trajectories from behavior in the baseline phase.

In addition to adaptation, we measured any potential compensatory behaviour produced in response to pitch perturbation, over and above changes driven by sensorimotor adaptation. For this analysis, we included only the sustained vowel sessions, as the short duration of the spoken words prevented measuring this compensatory behavior. Reflexive compensatory responses were measured in a window 200-300 ms after vocalization onset (chosen a priori) and 400-500 ms after onset (chosen after examining the data). For this analysis of compensatory behaviour, pitch trajectories (in cents) for each trial were normalized to the average value in that same trial during the 50-120 ms window after vocalization onset to isolate within-trial compensatory changes from any across-trial adaptive changes.

#### Experiments 2-4

Trajectories for each trial were transformed to a cents scale using the median *f0* value in a baseline window spanning the 50 ms window immediately following the onset of the perturbation. For Experiment 3, where the perturbation began immediately at the onset of vocalization, the *f0* value in a baseline window spanning from 50-120 ms after vocalization onset was calculated to transform the trajectories to cents. Omitting the first 50 ms avoids the frequent initial transient errors in pitch tracking in this period; note this window still avoids any potential compensatory behavior driven by the perturbed auditory feedback, which has a latency of ∼150 ms (Burnett et al., 1998). The median trajectory, in cents, was calculated across all unperturbed trials as a measure of participants’ typical baseline behavior in the absence of a perturbation (see Figure S1). This baseline trajectory was then subtracted from each perturbed trial to derive a normalized trajectory of compensatory behavior. Finally, these normalized compensatory trajectories in trials with an upwards pitch perturbation, expected to result in a downward response, were sign-flipped, such that compensation opposing the perturbation was reflected as a positive value in all trials.

### Statistical Analysis

For all studies, statistical testing was conducted in R (R Core Team, 2024) using linear mixed effects models (*LME4* package, Bates et al., 2015). Model selection was conducted using the *buildmer* package to select the best-fitting model (Voeten, 2023). Both forward and backward model selection procedures were applied to ensure robustness and consistency in model specification. Forward selection was based on likelihood ratio tests, starting with a simple model and sequentially adding predictors and random effects to identify the best-fitting model. For backward elimination, the procedure began with the full model, from which predictors were removed step by step until the most efficient model was reached. Both procedures used the default optimizer (*lmerControl*) to ensure stable model convergence. Using both approaches allowed cross-validation of the final model structure, minimizing the risk of overfitting or model misspecification that could arise from relying on a single selection path. Both forward and backward selection procedures yielded identical results in all cases; therefore, outcomes are reported based on the model obtained through forward selection. The final model was refitted using *lmer*, and statistical significance was assessed by maximum likelihood t-tests using the Satterthwaite’s method from the *lmerTest* package (Kuznetsova et al., 2017). The final fits were evaluated using type III ANOVAs and the *r.squaredGLMM* function from the *MuMIn* package (Bartoń, 2024).

For each fixed effect in the LMM, we report the F-value and corresponding p-value, as obtained from the linear mixed-effects analysis. Pairwise post-hoc comparisons report marginal means and standard errors. Effect sizes, where applicable, were reported as conditional R^2^ values for linear mixed-effects models (using *r.squaredGLMM*) and Cohen’s d for post-hoc comparisons and t-tests (using the *eff_size* from the package *EMMEANS* (Lenth, 2024). For all experiments, statistical significance was determined using α = 0.05.

The time course analysis of pitch trajectories during adaptation in words was conducted using Generalized Additive Mixed Models (GAMMs) built with the function *Bam* from the *mgcv* package (Wood, 2003, 2004, 2011, 2017; Wood et al., 2016) and the *itsadug* package (van Rij et al., 2022) for the figures. We included smooth terms for time by specifying a factor-smooth interaction for each experimental fixed effect (either group or condition), implemented as a thin plate regression with 10 basis splines. To account for inter-speaker variability in trajectories, we included a random smooth for each speaker over time, using a factor-smooth interaction spline with 10 basis functions and a first-order difference penalty. Results from the GAMM analysis were corrected for multiple comparisons with FDR with the *p.adjust* function in R (two separate comparisons for NH and CA groups comparing changes across phases and three comparisons testing for differences between NH and CA groups in the early exposure, late exposure and washout phases). Residual autocorrelation was modeled using an AR(1) process. For the within-group analyses, “Condition” was coded using treatment contrasts, with the baseline condition specified as the reference level.

#### Experiment 1

Experiment 1 included analyses from both adaptation and online compensation during the adaptation task. The initial models for the LMM analysis of sensorimotor adaptation included the fixed effects “phase” (baseline, early exposure, late exposure and washout) and “group” (CA versus NH) and their interactions, as well as random intercepts for individual speakers. Simple contrast coding was used such that the baseline phase served as the reference against which the comparisons were made. The baseline phase consisted of the 10 final baseline trials, the early exposure phase consisted of the first 10 trials of the exposure phase, late exposure phase included the final 10 trials of the exposure phase, and the washout phase consisted of the first three trials after the perturbation was removed. The early exposure phase was included given results from previous studies indicating that 1) maximum adaptation in pitch studies is typically reached quickly (see e.g., Alemi et al., 2020) and 2) adaptation may, in some cases, diminish over extended exposure to a perturbation (Alemi et al., 2020). Using 10 trials in each half of the exposure phase allowed us to account for the high variability in *f*0 from trial to trial. Conversely, adaptation in pitch has been shown to disappear rapidly (Alemi et al., 2020); using only the first three trials of the washout phase, while introducing some noise due to the small number of trials, is nonetheless necessary to measure this transient behavior.

For all five GAMM analyses (one for each group comparing across phases and three separate analyses comparing between groups in the early exposure, late exposure and washout phases) the model components were the dependent variable “pitch”, the fixed effect (“group” or “condition”), a smooth over time, and a random smooth over time by speaker. In the between group comparison, pitch trajectories were normalized by subtracting the mean trajectory in the baseline phase on a per-participant basis, yielding trajectories that represented changes from the baseline behavior.

Compensatory behaviour over and above adaptation during sustained vowel production was assessed by comparing pitch trajectories from the final 10 trials of the baseline phase,, to normalized trajectories (accounting for any adaptation effects) from the 40 trials produced in the exposure phase. Separate analyses were run for values extracted during an early window (200-300 ms after onset of the vocalisation) and late window (400-500 ms windows after vocalisation onset). The model included the fixed effects “phase” (baseline and exposure phase) and “group” (CA, NH), and the interaction between the two, as well as random intercepts for individual speakers.

#### Experiments 2-4

The same initial model structure was used for all three experiments, which included “group” (CA, NH), “direction of perturbation” (“up” and “down”) and their interactions, and random intercepts for individual speakers for an initial analysis. This analysis tested whether the two groups of participants differed from each other in their compensatory behaviour during pitch perturbation. Separate tests were conducted for early (200-300 ms after perturbation onset) and late (400-500 ms after perturbation onset) windows, similar to Experiment 1. An additional window was tested for Experiment 3, 400-500 ms after removing the perturbation, assessing whether participants fully returned to their unperturbed pitch trajectories. The length of the produced vowel was always longer than 1 second, allowing us to capture this window. To test whether CA and NH individuals showed compensation, we ran 4 additional one-sample t-tests against a normal distribution with 0 mean, one for each group and direction, using the *t-test* function in the *stats* package, included in R.

## Results

### Experiment 1

The goal of Experiment 1 was to examine whether CA participants, as a group, show reduced adaptation to perturbations of vocal pitch similar to the impaired adaptation observed in CA across all other motor domains that have been examined. Sustained vocalization is the standard behavior examined in studies of vocal motor control using perturbation paradigms with NH speakers and speakers diagnosed with neurodegenerative disorders (e.g., (Abur et al., 2018; Houde et al., 2019; Li et al., 2019). We thus included one assay of sustained vocalization of the vowel “ah” (/ɑ/) to compare our results with the existing literature on adaptation in NH individuals and individuals with neurological disorders other than ataxia. However, it is not clear what pitch target, if any, participants are using in these tasks. Vocal pitch tends to drift substantially over the course of multiple sustained vocalizations even in the absence of any perturbation (Alemi et al., 2020; 2021), suggesting that speakers may try to match the pitch they were producing at the end of one trial with the pitch at the start of the next trial. Moreover, the adaptive responses to pitch perturbations are larger when participants are instructed to reproduce a particular cued pitch compared to the more common instruction to “maintain a steady pitch (Jones & Keough, 2008; Keough et al., 2013), suggesting that speakers do not have a particular pitch target during sustained vocalization. Finally, sustained vocalization is a highly unnatural task, with limited ecological validity and the underlying control mechanisms might differ from pitch control during more typical speech production (Dahl et al., 2024). Thus, it is possible that changes in pitch behavior observed during repeated sustained vocalization reflect changes driven by other processes apart from sensorimotor adaptation, per se. To address these concerns, we additionally tested pitch adaptation during production of a simple spoken word (“bod”, /bɑd/).

#### Adaptation

In the sustained vowel production task (Figure 2A & B), both CA and NH speakers adapted to pitch perturbations during vowel production, shown by a main effect of phase (*F(*3,2748) = 10.69, *p* < .0001; *R²* = 0.24). While there was no overall difference between NH and CA participants, a significant interaction between group and phase was observed, consistent with our initial hypothesis of reduced adaptation in the CA group (*F*(3, 2748) = 2.80, *p* = 0.04).

NH speakers increased their pitch from baseline immediately following exposure to the perturbation (26 ± 5 cents, *t*(2753) = 4.69, *p* <.0001, *d = 0.31*) and maintained roughly the same level through the final 10 trials of the exposure phase (21± 5 cents, *t*(2753) = 3.88, *p* <.001, *d = 0.26*) and into the first three repetitions of the washout phase (25 ± 8 cents, *t*(2754) = 3.07, *p* = 0.01, *d = 0.30*).

The CA speakers showed a different pattern. Whereas the final 10 trials were produced at a higher pitch compared to the baseline (22 ± 6 cents, *t*(2754) = 3.75, *p* = 0.001, *d = 0.27*), similar to the NH group (NH-CA: 5±11 cents, *t*(133) = 0.43, *p* = 0.67), no significant adaptation was observed during the first 10 trials of the exposure phase in the CA group (8 ± 6 cents, *t*(2754) = 1.28, *p* = 0.58), unlike the immediate adaptation observed in the NH group (NH-CA: 24 ± 11 cents, *t*(133) = 2.102, *p* = 0.04, *d = 0.29*). Similarly, the changes observed at the end of the exposure phase in the CA group did not carry over into the washout phase, where CA participants showed no observable adaptation (7 ± 9 cents, *t*(2754) = 0.77, *p* = 0.87), again differing from the clear retention of adaptation seen in the NH group, though the direct comparison between groups failed to reach statistical significance (NH-CA: 24±14 cents, *t*(330) = 1.68, *p* = 0.09).

In the word production task (Figure 2C & D), we again found a main effect of phase, suggesting an overall presence of adaptation (*F*(3, 2726) = 8.73, *p* < .0001, *R²* = 0.29). As for the vowel production task, no main effect of the group was found, (*F*(1, 93) = 1.10, *p* = 0.30), but we did observe an interaction between group and phase (*F*(3, 2726) = 2.94, *p* = 0.03).

While the NH group showed an increase in pitch during the early part of the exposure phase compared to the baseline phase (17 ± 6 cents, *t*(2729) = 2.72, *p* = 0.03, *d* = 0.18), no such adaptive response was observed in the CA group (-9 ± 7 cents, *t*(2738) = -1.30, *p* = 0.57). There was a steady decline in pitch throughout the rest of the experiment observable in both groups. For the NH group, this resulted in a return of pitch to roughly the baseline level during the late exposure and washout phases (NH: *late exposure*: -0.5 ± 7 cents, *t*(2730) = -0.08, *p* = 1.00; *washout*: -17± 9 cents, *t*(2729) = -1.80, *p* = 0.27). For the CA group on the other hand, this same downward trend resulted in a lower pitch during the late exposure and washout phases than during the baseline phase (CA: *late exposure*: -21 ± 7 cents, *t*(2738) = -3.05, *p* =0.01, *d* = -0.22; *washout*: -37 ± 10 cents, *t*(2733) = -3.67, *p* < 0.01, *d* = -0.40). While the pitch was higher for the NH than the CA group across all phases after the baseline, likely driving the significant interaction between group and phase, direct comparisons between the NH and CA groups in each phase did not reach statistical significance (all *p* > 0.11).

Unlike the steady-state pitch produced in sustained vocalization, however, pitch production in isolated English words consists of a large decline in pitch over the course of the vowel (Figure 3A & B). As a follow-up test, we used GAMMs to compare the full pitch trajectories produced in the early exposure, late exposure, and washout phases to the baseline phase to ensure that the lack of adaptation we observed in the CA group was not due to our chosen time window failing to capture changes in some part of this trajectory.

**Figure 3:**
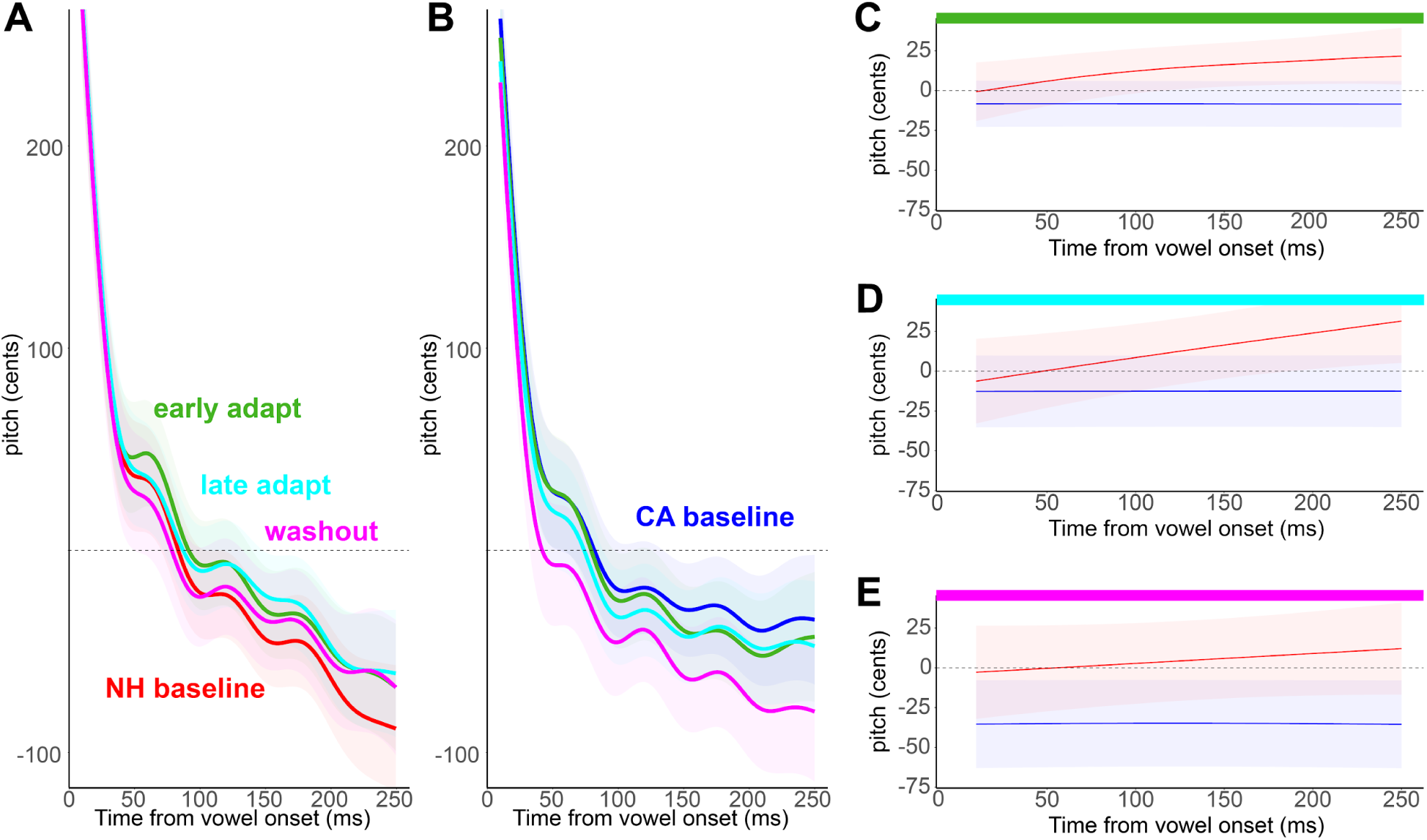
GAMM adaptation results for the word production task from Experiment 1. In all panels, red indicates values for NH speakers and blue indicates values for CA speakers. Shading shows standard error. **A**: Pitch trajectory smooths during word production for NH speakers in the baseline (red), early exposure (green), late exposure (cyan) and washout (magenta) phases. **B**: Pitch trajectory smooths during word production for CA speakers in the baseline (blue), early exposure (green), late exposure (cyan) and washout (magenta) phases. **C**: Pitch trajectory smooths in the early exposure phase, normalized by subtracting the average trajectory in the baseline phase. **D-E**: As for C, showing the late exposure and washout phases, respectively.

The results of these models largely replicated the window-based approach. NH speakers (Figure 3A, 56.8% variance explained by the model) showed increased pitch consistent with adaptive responses in both the early exposure phase (*β* = 13.73, *SE* = 2.75, *t* = 4.98, *p* < 0.0001, adjusted *p* = 0.0001) and late exposure phase (*β* = 14.24, *SE* = 2.77, *t* = 4.14, *p* < 0.0001, adjusted *p* = 0.0001), with pitch returning to close to the baseline trajectory in the washout phase (*β* = 4.88, *SE* = 4.06, *t* = 1.20, *p* = 0.23, adjusted p = 0.23). Conversely, the CA speakers (Figure 3B, 48.8% variance explained) did not show any increase in pitch. Rather, they showed a slightly reduced pitch during the early exposure phase (*β* = –7.39, *SE* = 3.35*,t* = –2.21, *p* = .03, adjusted *p* = 0.04), after which pitch continued to decline through the late exposure phase (*β* = –11.95, *SE* = 3.34, *t* = –3.58, *p* < .001, adjusted *p* < .001) and washout phase (*β* = –33.95, *SE* = 4.94, *t* = –6.87, *p* < .001, adjusted *p* < .001). As expected given the overall decline in pitch throughout the vowel, significant nonlinear effects of time were observed across all conditions for both speaker groups; random smooths for speakers improved the fits for both NH and CA models (Table S1).

Directly comparing the CA and NH groups (Figure 3C-E) showed that the NH speakers produced significantly higher pitch trajectories than the CA speakers during the early exposure phase (28.8% deviance explained, *β* = 21.88, *SE* = 9.88, *t* = 2.21, *p* = .03; adjusted *p* = .04) and washout phase (69.5% deviance explained, *β* = 39.82, *SE* = 18.82, *t* = 2.12, *p* = 0.03; adjusted *p* = 0.04). While the same trend for higher pitch in the NH group was observed in the late exposure phase, the direct comparison in this phase did not reach statistical significance (49.2% deviance explained, *β* = 25.87, *SE* = 15.51, *t* = 1.67, *p* = 0.09, adjusted *p* = 0.11). Analysis of the smooth terms indicated significant fluctuations over time in both groups, though with different degrees of complexity. During the early and late exposure phase, only that NH group exhibited significant changes over time, with higher pitches (relative to baseline) later in the trajectory. During the washout phase, both the ataxia and the healthy group exhibited significant time-dependent changes, though only for the NH group was the trend clearly to increase pitch (relative to baseline) over the course of the trajectory. These non-linear effects suggest that adaptation in pitch production affects the later, lower components of the trajectory rather than the initial high pitch, at least for this type of pitch contour in English. For all three phases, the random smooth for participants captured substantial inter-individual variability.

Together, these results suggest that NH speakers adapted their pitch in word production in response to the auditory perturbation, observable most visibly in the early phase of the exposure period. Conversely, the CA group showed no adaptive increase in pitch, showing instead a steady decrease in pitch that continued to grow over the course of the experiment. While the pitch for both groups declined from the early adaptation phase through the washout, the pitch in the NH group remained consistently above that in the CA group. This suggests that, in the NH group, adaptation here served to counteract a natural downward trend in pitch over the course of the experiment (see Discussion).

#### Compensation over and above adaptation

Investigating compensatory responses later in the trial that may occur over and above any trial-to-trial changes (Raharjo et al., 2021), we assessed whether the previous findings of enhanced compensation in individuals with CA as a response to inconsistent, transiently induced perturbations, can be generalized to compensatory responses when consistently exposed to perturbations beginning at vocalization onset.

To explore the existence of this potential behaviour, we first examined compensatory responses that occur over and above adaptive changes to consistent perturbations during the window from 200-300 ms after vocalisation onset (Figure 4). Responses in this early window were significantly larger in the direction opposing the perturbation than in the baseline phase across both groups (18 ± 3 cents, *F*(1,4114) = 26.93, *p* < .0001; *R²* = 0.20), confirming the presence of online compensatory behavior in addition to any trial-to-trial changes in production. No difference was found between the CA (16.06 ± 7.16 cents) and NH (19.95 ± 7.71 cents) groups (*F*(1, 96) = 2.22, *p* = 0.14), suggesting the magnitude of this compensatory response was intact, but not enhanced, in the CA group. No interactions were observed (*F*(1, 4114) = 0.45, *p* = 0.50).

**Figure 4:**
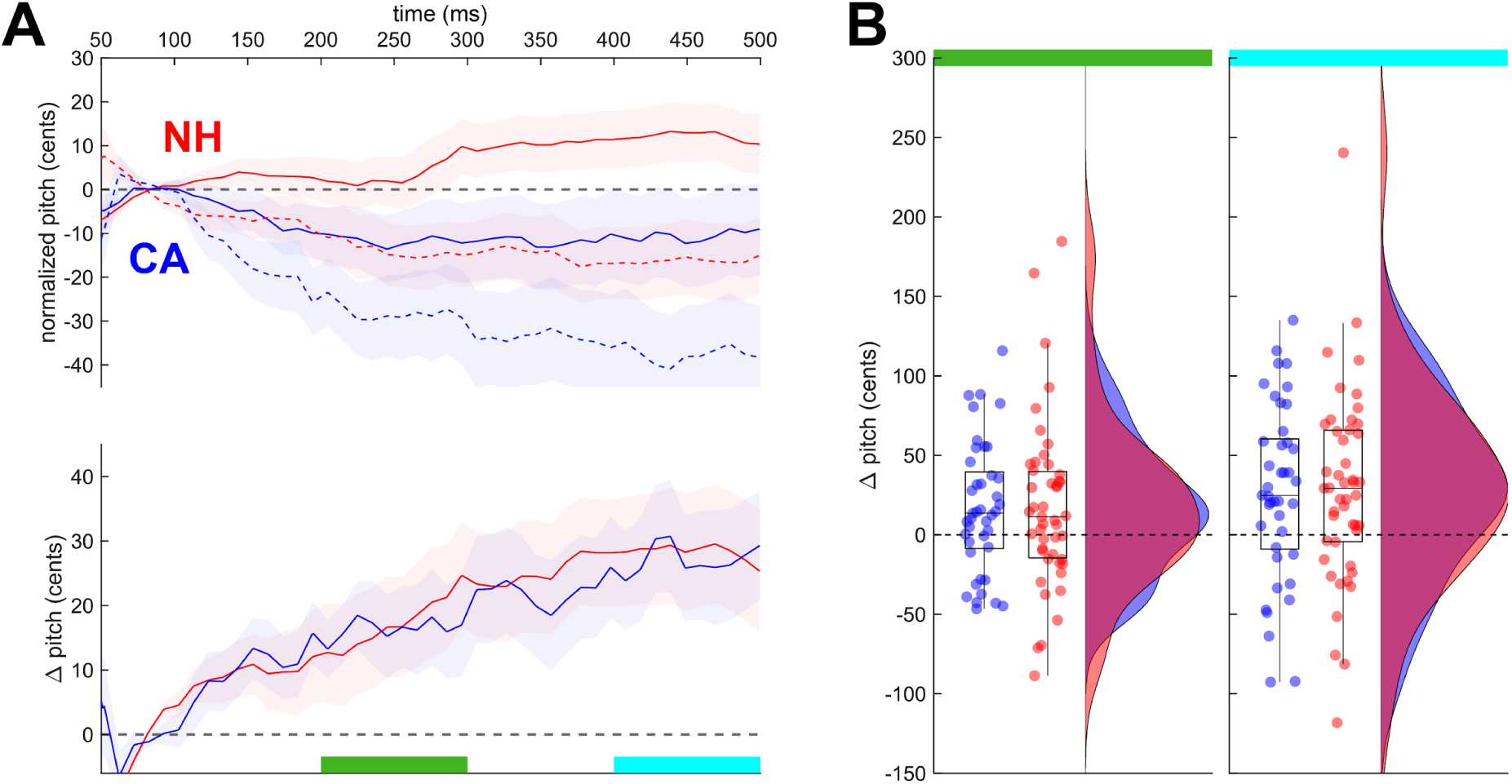
Compensation during exposure to consistent auditory perturbations during vowel production, over and above any adaptive changes in behavior. In all panels, blue indicates values for CA speakers and red indicates values for NH speakers. **A**: Top: mean pitch trajectories from the perturbed session during the exposure phase (solid lines) and the final 10 trials of the baseline phase (dashed lines). All trajectories were normalized to the unperturbed session. Shading shows standard error across participants. Bottom: online compensation, shown by the difference between pitch trajectories during the exposure phase and the mean pitch trajectory during the final 10 trials of the baseline phase. Shading shows standard error across participants. Green and cyan highlighting show the early (200-300 ms post onset) and late (400-500 ms post onset) analysis windows, respectively. **B**: Individual participant behavior in the early (left) and late (right) analysis windows. Dots and density plots show individual participant mean responses to each perturbation direction (up/down, 2 points per participant).

After examining the data, we observed that compensatory pitch responses had not reached their plateau in our planned analysis window, but for both groups did plateau roughly 400 ms after vocalization onset (Figure 4A), suggesting our planned window likely failed to capture the true extent of compensation. Subsequently, we conducted a post-hoc analysis using a later window capturing the maximal plateau of compensation, 400-500 ms after vocalization onset. Again, both groups produced a significantly higher pitch during the exposure phase than the baseline phase (difference between the phases across groups: 30 ± 4 cents, (*F*(1, 4108) = 54.53, *p* < .0001, *R²* = 0.24), but again, the CA and NH groups did not differ in their responses (*F*(1, 95) = 3.13, *p* = 0.08) and no interactions were observed (*F*(1, 4108) = 0.39, *p* = 0.53).

### Experiment 2

Previous studies have shown that, compared to NH speakers, individuals with CA produced larger compensatory responses to unexpectedly introduced pitch perturbations during ongoing vocalisations, potentially reflecting increased reliance on feedback control strategies in the presence of impaired feedforward control (Houde et al., 2019; Li et al., 2019). To explore whether compensatory behaviour to pitch perturbations in the current group of individuals with CA showed similar patterns, we examined compensatory changes in *f0* to unexpected vocal pitch perturbations which began 200-500 ms after vowel onset, and which varied in direction across trials. As for the compensation results in Experiment 1, we report results from both a planned “early” window 200-300 ms after perturbation onset and a “late” window 400-500 ms after vocalization onset that better captures the maximum compensatory responses.

In the early window (Figure 5), both CA and NH groups showed opposing compensatory responses to both perturbation directions (all *p* < .001, d ≥ 0.54, see Table S2). Upward perturbations resulted in slightly smaller responses than downward perturbations across both groups (Up-Down: -4 ± 1 cents, *F(*1,6931) = 11.20, *p* < .001; *R²* = 0.07). NH and CA speakers did not show differences in the magnitude of compensation (*F(*1, 88) = 2.53, *p* = 0.12), nor was there any interaction observed between group and perturbation direction (*F*(1, 6931) = 0.97, *p* = 0.32).

**Figure 5:**
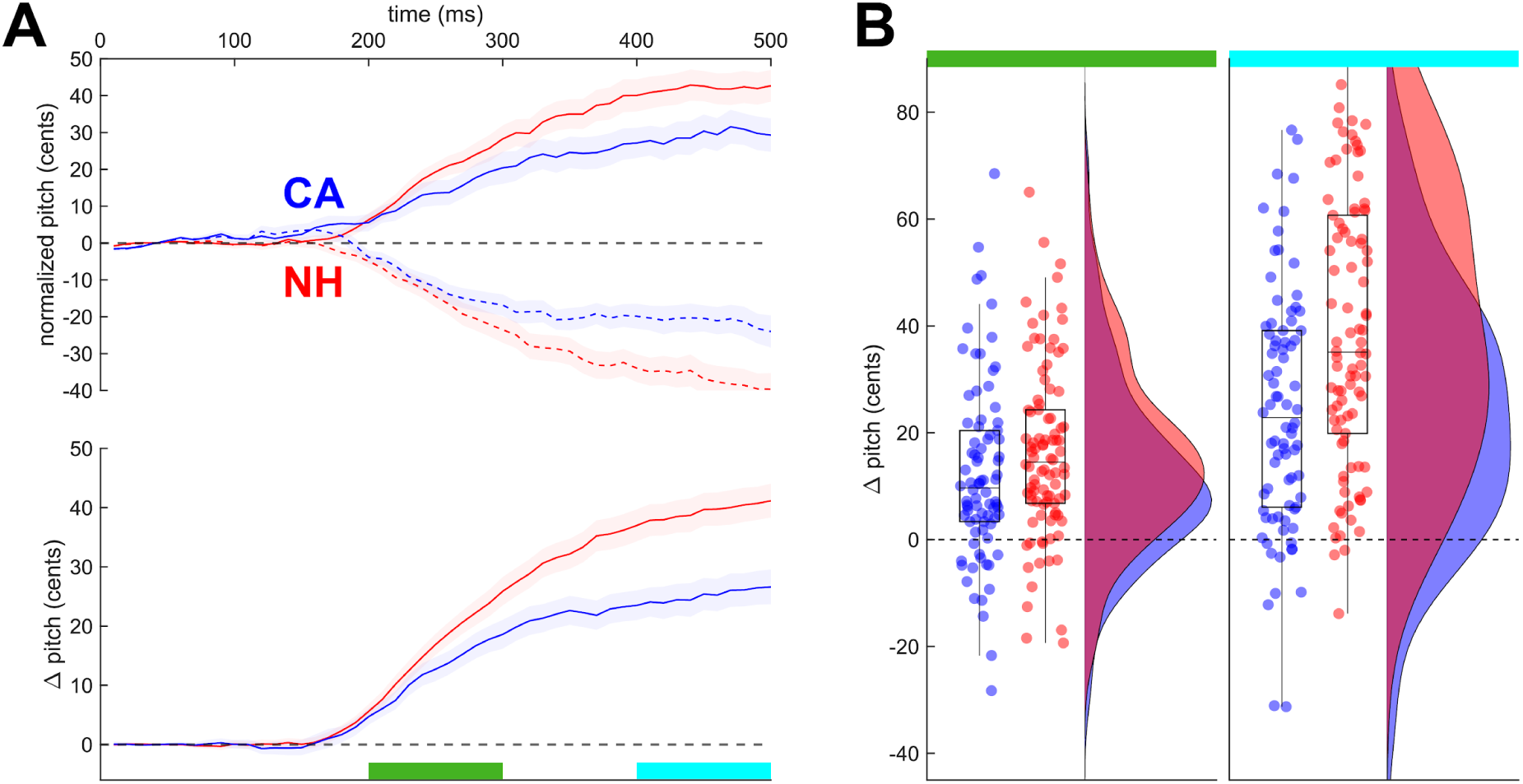
Compensation during sustained auditory perturbations delivered mid-vowel. In all panels, blue indicates values for CA speakers and red indicates values for NH speakers. **A**: Top: online compensation to upward (solid lines) and downward (dashed lines) perturbations, normalized by unperturbed trials. Shading shows standard error across participants. Bottom: compensation across both perturbation directions, measured by the mean pitch trajectory combining responses to upward (sign-flipped) and downward perturbations. Shading shows standard error across participants. Green and cyan highlighting show the early (200-300 ms after perturbation onset) and late analysis (400-500 ms after perturbation onset) windows, respectively. **B**: individual participant behavior in the early (left) and late (right) analysis windows. Dots and density plots show individual participant mean responses to each perturbation direction (up/down, 2 points per participant).

Post-hoc examination during a later time window from 400-500 ms after vowel onset, however, revealed a different pattern of results. While both CA and NH groups showed a significant compensatory response to both perturbation directions (all *p* < .001, all *d* ≥ 0.83, see Table III, Supplementary materials), individuals with CA showed smaller compensatory responses in this later time window than NH speakers (NH-CA: 14 ± 5 cents, *F*(1,88)= 8.98, *p* < 0.01; *R²* = 0.10). Responses to the upward perturbation in this late window were again smaller than for the downward perturbation for both groups (-8 ± 1 cents, *F*(1, 6902) = 27.78, *p* < .0001). No interactions between the group and direction of perturbation were observed (*F*(1, 6902) = 1.21, *p* = 0.28).

### Experiment 3

Results assessing compensation from Experiments 1 and 2 show diverging results. While no group difference between the CA and NH groups was found in compensation to pitch perturbations when they are applied continuously and consistently (Experiment 1), we found slightly reduced compensation in the CA group to inconsistent and unpredictable pitch perturbations delivered mid-vocalization (Experiment 2), especially when examining the maximal response 400-500 ms after vocalization onset. It is possible that this difference may have arisen either from differences in the consistency of perturbation across trials (predictable vs unpredictable) or from the timing of the perturbation (vocalization onset vs. mid-vocalization). In support of the latter, previous work has suggested that compensatory responses to perturbations beginning at the onset of vocalization may rely on a different mechanism than compensating to mid-vowel perturbations: compensatory responses to onset perturbations must necessarily compare the auditory feedback signal to a desired target or predicted state, whereas responses to perturbations later in the vocalisations may be controlled by comparing reafferent auditory feedback information to the previous (ongoing) *f0* production to maintain a steady pitch (Hawco & Jones, 2009; Larson et al., 2001; Scheerer & Jones, 2018). After a preliminary (numerical) examination of data from 12 initial participants in both the CA and NH groups, which hinted at the pattern eventually observed in the full dataset, we added an additional study to the larger battery of experiments to examine compensatory responses to unpredictable perturbations (which varied in direction across trials) delivered at vocalization onset, allowing us to test whether the timing of the perturbation was a source of the difference in compensatory behavior of the CA group between Experiments 1 and 2.

During the early window, 200-300 ms after onset, the CA group showed statistically significant evidence of a compensatory response to the upward perturbation (*p* = 0.05, *d* = 0.38) and NH speakers showed a significant response to the downward perturbation (*p* = 0.03, *d* = 0.44). The remainder of the responses were not significant (*p* ≥ 0.57, Figure 6 and Table S3). CA and NH did not differ in their responses to onset perturbations (*F*(1, 56) = 0.23, *p* = 0.63; *R²* = 0.005) and responses to upwards perturbations did not differ from responses to downward perturbations (*F*(1, 4422) = 0.57, *p* = 0.45). Group and condition, however, did interact (*F*(1, 4422) = 11.27, *p* < .001). This interaction was caused by the fact that CA speakers differed in their responses to down- and upward perturbations, with downward perturbations showing less compensatory response than upward perturbations (Up-Down: 9 ± 3 cents, *z*(inf) = 2.96, *p* = .01). This pattern also caused a difference between NH and CA speakers in the sense that CA speakers compensated less than NH for downward perturbations (CA-NH: -9 ± 3 cents, *z*(inf) = -2.60, *p* = 0.04). The two groups did not differ in response to upward perturbations (CA-NH: 6 ± 3 cents, z(inf) = 1.88, *p* = 0.24) and NH speakers did not show a difference between down and upward perturbations (Down-Up: 6 ± 3 cents, *z*(inf) = 1.81, *p* = 0.28).

**Figure 6:**
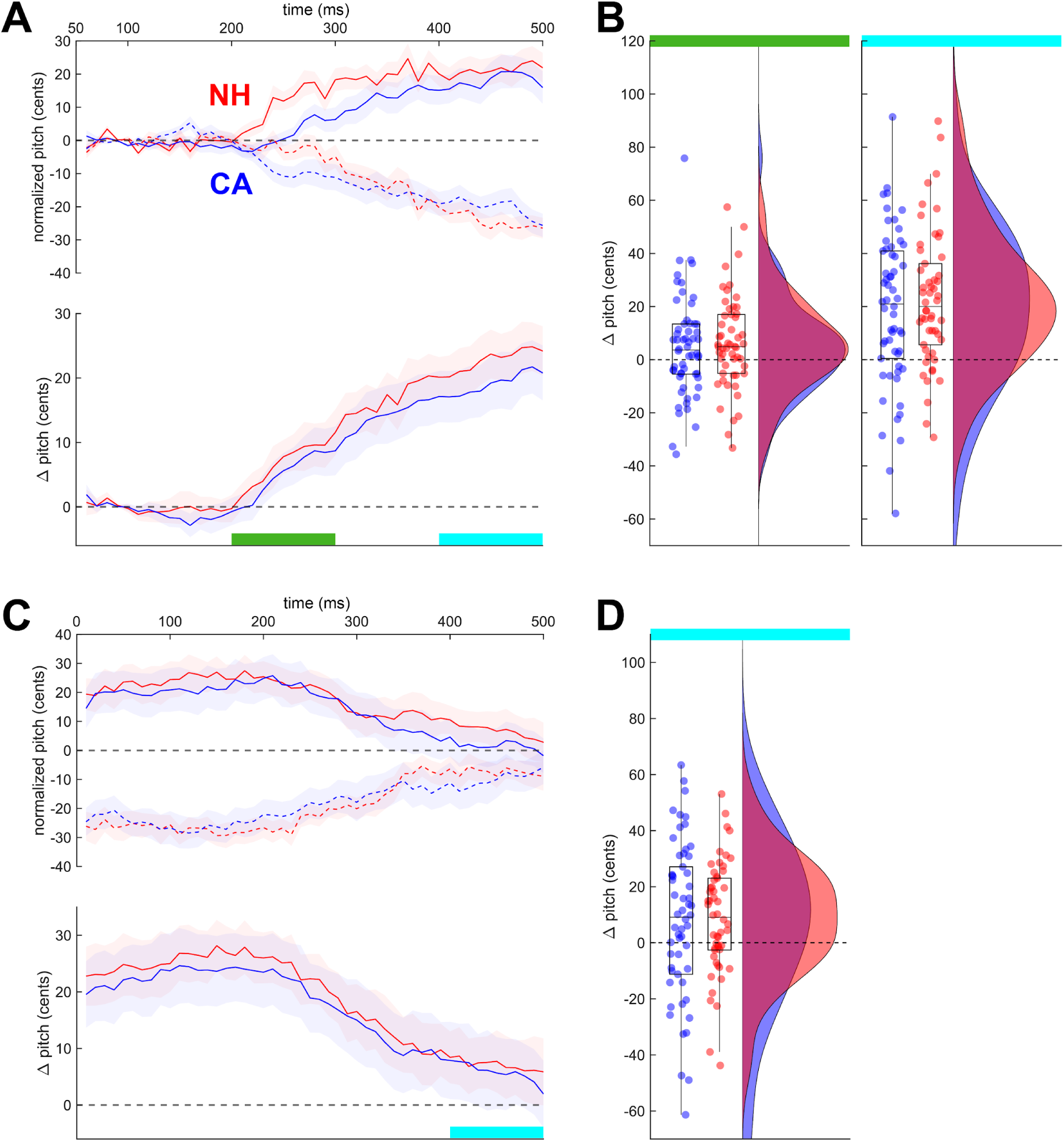
Compensation to transient auditory perturbations delivered at vowel onset. In all panels, blue indicates values for CA speakers and red indicates values for NH speakers. **A:** Top: online compensation to upward (solid lines) and downward (dashed lines) perturbations, normalized by unperturbed trials. Shading shows standard error across participants. Bottom: compensation across both perturbation directions, measured by the mean pitch trajectory combining responses to sign-flipped upward and downward perturbations. Shading shows standard error across participants. Green and cyan highlighting show the early (200-300 ms after vowel/perturbation onset) and late (400-500 ms after vowel/perturbation onset) analysis windows, respectively. **B**: Individual participant behavior in the early (left) and late (right) analysis windows. Dots and density plots show individual participant mean responses to each perturbation direction (up/down, 2 points per participant). **C:** As for A, showing the pitch response after the removal of the perturbation (time 0 = perturbation offset). **D**: As for B, showing pitch response 400-500 ms after perturbation removal.

As for Experiment 2, we also inspected the compensatory behaviour during a 400-500 ms window after perturbation (and vocalization) onset to estimate the maximum magnitude of this response. Although the window includes, in some cases, up to 100 ms after the perturbation was removed (which occurred randomly between 400 to 550 ms after vocalization onset), it is nonetheless possible to measure compensation in this window as the behavioral latency to auditory feedback is ∼150 ms (e.g., Burnett et al., 1998). In this later window, compensation was clearly visible and significant for both CA and NH for both perturbation directions (all *p* ≤ 0.03, all d ≥ 0.44, Table S3). The magnitude of the response did not differ between the CA and NH groups (*F*(1, 56) = 0.51, *p* = 0.48; *R²* = 0.03) nor between the upward and downward perturbations (*F*(1, 4418) = 3.26, *p* = 0.07), nor was there any interaction between these factors (*F*(1, 4418) = 0.90, *p* = 0.34).

As an additional analysis of feedback-driven compensation, we analyzed the period following the removal of the pitch perturbation (starting 400-550 ms after vocalization, across trials; Figure 6C&D). Our reasoning here is that the removal of the pitch alteration can itself act as a type of perturbation, since speakers have begun to alter their behavior to correct for the initial alteration. Pitch tracks were aligned to the perturbation offset, and we examined a window 400-500 ms following this time point, expecting that the produced pitch would return to baseline values. This was indeed the case: for both groups, the produced pitch in this window, though numerically slightly above baseline, did not differ from 0 for either perturbation direction (*p* ≥ 0.13, d ≤ 0.30, Table S3). No differences in behaviour were found between CA and NH groups (*F*(1,55) = 1.16, *p* = 0.29; *R²* = 0.01) or up or downward perturbations (*F*(1,4371) = 0.85, *p* = 0.36), nor was there an interaction between these factors (*F*(1, 4372) = 2.17, *p* = 0.14).

### Experiment 4

Experiment 3 was designed to mediate between alternative explanations for the different results in compensation between Experiments 1 and 2: perturbation consistency or perturbation timing. The results, which largely mirror Experiment 1, suggest that the reduced compensatory response in the CA group in the late window in Experiment 2 may stem from the timing of the perturbation starting mid-vocalization versus at vocalization onset. This result emphasizes that changes in pitch perturbation timing may have unanticipated effects on the resulting compensatory behavior. Experiment 4 was designed to address a separate aspect of perturbation timing that may have affected the results in Experiment 2. Our initial (numerical) inspection of preliminary results from a subset of participants in Experiment 2, which was consistent with the final result, suggested equal or reduced compensation in the CA group compared to the NH group, a striking difference from previous studies showing increased pitch compensation in individuals with CA (Houde et al., 2019; Li et al., 2019). However, previous studies used transient perturbations that began after vocalization onset and, perhaps crucially, were removed before vocalization offset, differing from the perturbations in Experiment 2 which remained in place until the end of the trial. Experiment 4 was designed to test whether the seemingly minor difference in perturbation offset timing may have driven this difference in compensatory behavior. Here, the perturbation was randomly introduced, but removed after 400 ms. Note that as for Experiment 3, the late analysis window 400-500 ms after perturbation onset, while occurring after the removal of the perturbation in real time, nonetheless captures compensatory behavior to the perturbation given the >100 ms latency of auditory feedback responses.

In the early window 200-300 ms after perturbation onset, both groups of speakers compensated for both perturbation directions (all *p* ≤ 0.04, d ≥ 0.76, Figure 7 and Table S4). There was no difference in the magnitude of compensation between the NH and CA groups (*F*(1, 32) = 0.05, *p* = 0.82; *R²* = 0.05), no difference between upward and downward perturbations *(F*(1, 2525) = 0.16, *p* = 0.69) nor an interaction between these factors (*F*(1, 2525) = 1.23, *p* = 0.27).

**Figure 7:**
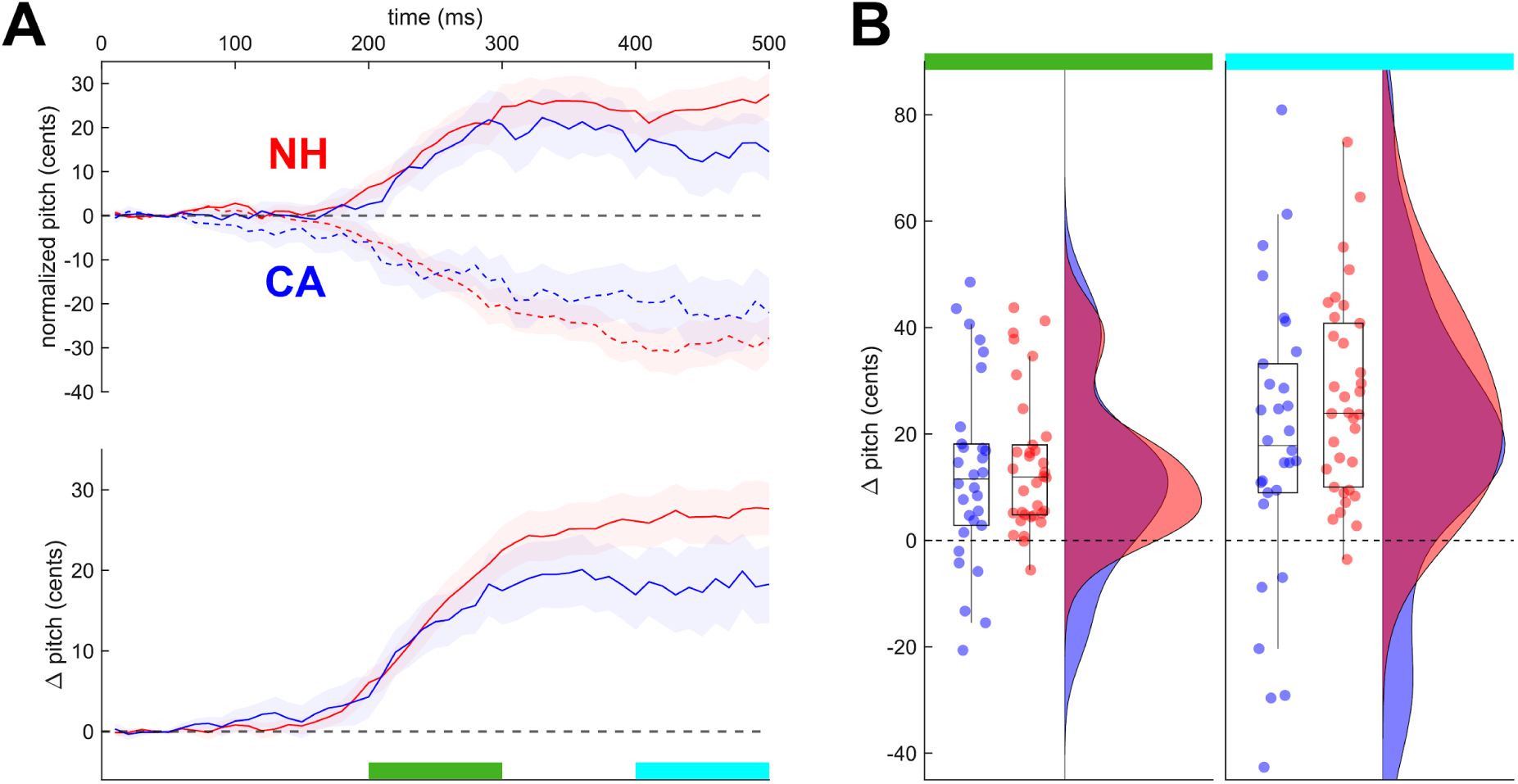
Compensation to transient auditory perturbations delivered mid-vowel. In all panels, blue indicates values for CA speakers and red indicates values for NH speakers. **A:** Top: online compensation to upward (solid lines) and downward (dashed lines) perturbations, normalized by unperturbed trials. Shading shows standard error across participants. Bottom: compensation across both perturbation directions, measured by the mean pitch trajectory combining responses to sign-flipped upward and downward perturbations. Shading shows standard error across participants. Green and cyan highlighting show the early (200-300 ms after perturbation onset) and late (200-300 ms after perturbation onset) analysis windows, respectively. **C**: Individual participant behavior in the early (left) and late (right) analysis windows. Dots and density plots show individual participant mean responses to each perturbation direction (up/down, 2 points per participant).

In the late window 400-500 ms after perturbation onset, apart from the CA speakers, who showed only a trend as a response to downward perturbation (*p* = 0.06, *d* = 0.54), both CA and NH showed large compensatory responses to both upward and downward perturbations (*p* ≤ 0.05, *d* ≥ 0.54, Table S4). While the results numerically resembled Experiment 2, with the CA group showing a slightly reduced compensation magnitude compared to the NH group, this between-group difference was much smaller and, crucially, was not statistically significant (*F*(1, 32) = 2.62, *p* = 0.12). While upward perturbations resulted in larger compensatory responses than downward perturbations (Up-Down: 5 ± 2 cents, *F*(1, 2520) = 5.16, *p* = 0.02; *R²* = 0.07), this effect was similar for both groups as shown by the lack of any interaction effect (*F*(1, 2520) = 0.43, *p* = 0.51).

## Discussion

Sensorimotor adaptation has been shown to be impaired in individuals with CA across limb movements, saccades, walking, and oral articulation, indicating a critical, domain-general role of the cerebellum in this type of motor learning. We investigated whether the control of pitch, which differs substantially from other motor control activities in its higher reliance on feedback control than predictive/feedforward control, shows similar impairments in sensorimotor adaptation in individuals with CA. When we examined sensorimotor adaptation of pitch during both word production (a natural speech task) and during sustained phonation (the standard task in measuring pitch adaptation), we found that NH speakers produced an immediate adaptive response at the onset of the applied pitch perturbation while the CA group showed no change in behavior, actually showing a slight decrease in pitch over the course of the experiment in the word production task. Thus, despite the differences in reliance on predictive control in pitch compared to reaching and oral articulation, our results suggest the cerebellum likely plays a similar role in sensorimotor adaptation of vocal pitch as in other motor domains. Notably, the absence of any observable adaptation in the CA group, compared to the present but reduced adaptation observed in this group in other motor domains, may suggest that the cerebellum is particularly critical for regulation of vocal pitch.

While the initial patterns of adaptation (and lack of adaptation in the CA group) in the early exposure phase were similar in both word production and sustained phonation, behavior in the two tasks diverged in the later phases of the experiment. In the sustained phonation task, the NH group showed an immediate adaptive response at the very start of the exposure phase, and maintained this elevated pitch through the late exposure phase and into the first few trials of the washout phase, as in previous studies (Alemi et al., 2020; Behroozmand & Sangtian, 2018). Such aftereffects are a hallmark of sensorimotor adaptation across motor domains. Additionally, while adaptation is typically gradual in other motor domains, adaptation in pitch has been shown to be very rapid (Alemi et al., 2020; Behroozmand & Sangtian, 2018; Jones & Keough, 2008; Keough et al., 2013), consistent with the observed behavior in the NH group. Conversely, while the CA group produced an elevated pitch during the late exposure phase, they showed no evidence of adaptation in the early exposure phase, nor in the washout phase, returning immediately to baseline behavior with no visible aftereffects. The lack of initial adaptation and any aftereffects, given the presence of learning in these phases both in the NH data and in the broader literature on pitch adaptation, strongly suggest that adaptation did not occur in the CA group. However, the mechanism driving elevated pitch production in the late exposure phase in this group remains unclear. Importantly, the task participants perform in this paradigm is vaguely defined; although we followed typical instructions that participants “try to maintain a steady pitch”, it’s unlikely that participants are attempting to maintain some target, absolute pitch. This is shown by the gradual increase of pitch over time in the control session, when no perturbation is given. One possibility is that speakers try to reproduce the pitch they were producing on the previous trial; because the CA speakers continue to produce compensatory responses that oppose the perturbation, this self-imitation may drive increases in pitch over time, resulting in behavior at the end of the exposure phase that is superficially similar to adaptation-driven changes.

In the spoken word task, the NH participants immediately adapted to the perturbation (again, consistent with expectations from the pitch adaptation literature), but showed a subsequent downward trend in pitch, such that no aftereffects were observed in the washout phase. Notably, the CA group produced a similar downward trend, with pitch immediately lower than baseline in the early exposure phase that continued to drift downward over the course of the experiment. This consistent downward trend in both groups suggests that all participants showed a trend to lower their pitch over the course of the experiment, and that the adaptation visible in the NH group occurred on top of this trend; crucially, in all phases, the NH group produced higher pitch values than the CA group. This downward trend is surprising, given that no similar trend was observed in the control session for this task. However, the perturbation session always preceded the control session; it is likely that this downward trend reached a plateau by the start of the second session.

While the main aim of the current study was to assess cerebellar contributions to adaptation of vocal pitch, we also measured feedback-driven, real-time compensation to pitch perturbations. Unexpectedly, we found that compensatory responses of speakers with CA resembled the responses by neurobiologically healthy speakers and, as such, we did not find evidence for the expected over-compensation previously reported in this population (Houde et al., 2019; Li et al., 2019) in our cohort of speakers. By including several additional studies with slightly different perturbation paradigms, we excluded the possibility that the timing of the perturbation within a trial contributed to our divergent results. Several additional factors, though, might have contributed to the lack of overcompensation in our study. First, our cohort of speakers differs from the previous studies. Li et al. (2019) included 1 speaker with SCA1 and 11 speakers diagnosed with SCA3; our study contained a more diverse set of SCA types and it is yet unknown whether differences in type of cerebellar ataxia affect compensatory behaviour. However, Houde et al. (2019) showed a similar overcompensation in a more diverse group of SCA types (2 SCA2, 2 SCA3, 1 SCA5, 2 SCA6, 1 SCA7, 2 SCA8, 6 idiopathic cerebellar atrophy), making this explanation unlikely. Second, the current study had a much larger sample size (45 individuals with CA) compared to 12 in Li et al. (2019) and 16 in Houde et al.(2019). If indeed increased feedback-based responses are an idiopathic response to inaccurate feedforward control (Houde et al., 2019), the presence of some individuals with such an increased response may bias group results more in small samples. Third, although we controlled for the timing of the perturbation within the trial, there were still several methodological differences between these studies. Li et al. (2019) had participants produce long, sustained vocalizations, with several perturbations (±200 or ±500 cents) applied briefly in each vocalization. Houde et al. used a design more similar to that in our Experiment 4, with a single transient ±100 cent perturbation, though with no unperturbed trials. In both cases, the constant presence of pitch perturbations may have induced the CA group to switch to a more feedback-driven control mode in response to the unstable feedback, while the presence of unperturbed trials in our experiments (or, in the case of Experiment 1, the consistently applied perturbation) may have prevented such a switch.

In summary, our results show that sensorimotor adaptation to externally-applied perturbations of vocal pitch is severely impaired in individuals with cerebellar ataxia, strongly supporting a critical role of the cerebellum in this process. This is consistent with a domain-general role for the cerebellum in adapting motor behavior in response to sensory errors. The complete absence of adaptation observed in the CA group differs from the impaired, but present, adaptation in this group typically observed in other motor domains, suggesting that the cerebellum may be particularly critical for the adaptive control of vocal pitch. Conversely, we found that online, feedback-driven compensation to pitch perturbations was intact in the CA group. Although this is similar to the observed behavior in other motor domains, this differs substantially from the enhanced compensatory response previously demonstrated in this population. This result suggests that any enhanced reliance on feedback for pitch control is not an automatic result of cerebellar damage, but may rather be an individual-specific learned response, perhaps due to instability in the predictive control of pitch.

## Acknowledgements

This work was supported by grants from the National Institute on Deafness and Other Communication Disorders (Grant R01 DC017091), a core grant to the Waisman Center from the National Institute of Child Health and Human Development (Grant P50 HD105353), and an NIH High-End Instrumentation grant (S10 OD030415). We would like to thank Dr. Elaine Kearney for her support in developing the initial code for pitch perturbation with Audapter, Chris Naber for his technical support, and Dr. Yuyu Zeng for her advice on Generalized Additive Modelling.

## Supplementary material

**Figure S1.**
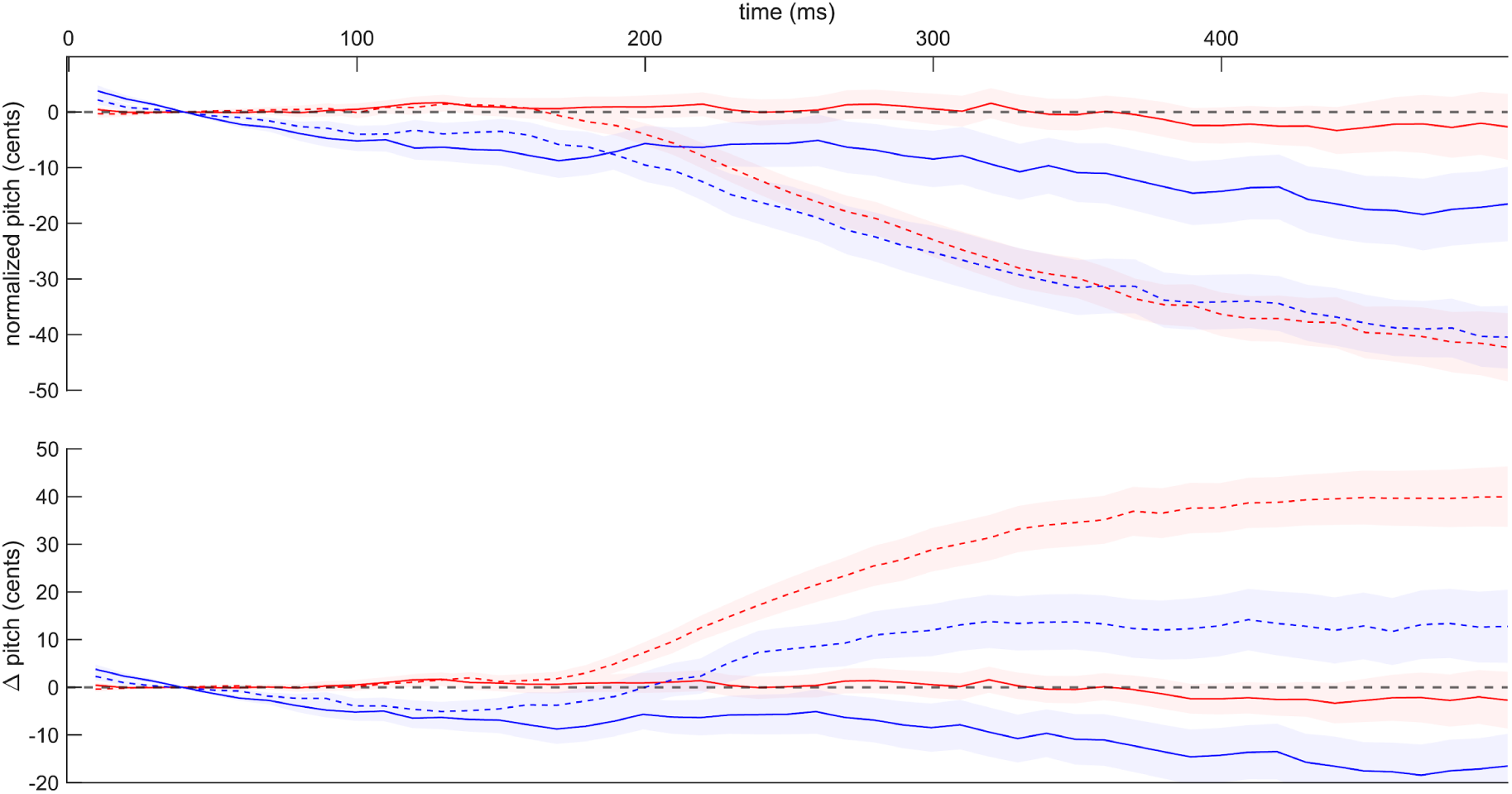
Compensatory responses for unperturbed and perturbed trials. Top panel: Dashed lines: mean pitch response to upward perturbations; solid lines: mean pitch trajectories during unperturbed trials. Shaded areas represent standard error. Blue: CA; red: NH. **Bottom panel:** As for the top panel, showing response to downward perturbations (dashed lines).

**Table S1:** Results for individual smooth terms from the Generalized Additive Mixed Models.

| CA |  |  |  |  |
| --- | --- | --- | --- | --- |
|  | *edf* | *Ref.df* | *F* | *p* |
| s(Time):baseline | 8.777 | 9 | 97.6 | **<2e-16** |
| s(Time):early exposure | 8.786 | 9 | 92.71 | **<2e-16** |
| s(Time):late exposure | 8.772 | 9 | 91.44 | **<2e-16** |
| s(Time):washout | 8.712 | 9 | 58.69 | **<2e-16** |
| s(Time,Speaker) | 374.58 | 409 | 21.79 | **<2e-16** |
| NH |  |  |  |  |
| s(Time):baseline | 8.801 | 9 | 97.62 | **<2e-16** |
| s(Time):early exposure | 8.83 | 9 | 98.31 | **<2e-16** |
| s(Time):late exposure | 8.779 | 9 | 97.82 | **<2e-16** |
| s(Time):washout | 8.691 | 9 | 56.07 | **<2e-16** |
| s(Time,Speaker) | 436.446 | 469 | 29.34 | **<2e-16** |
| early exposure |  |  |  |  |
| s(Time):CA speakers | 0.06856 | 9 | 0.003 | 0.252 |
| s(Time):NH speakers | 1.65456 | 9 | 0.568 | **2.59e-07** |
| s(Time,Speaker) | 504.66896 | 868 | 2.649 | **<2e-16** |
| late exposure |  |  |  |  |
| s(Time):CA speakers | 0.0367 | 9 | 0.001 | 0.0874 |
| s(Time):NH speakers | 0.9648 | 9 | 0.593 | **<2e-16** |
| s(Time,Speaker) | 629.9296 | 868 | 6.772 | **<2e-16** |
| washout |  |  |  |  |
| s(Time):CA speakers | 0.09425 | 9 | 0.009 | **0.0134** |
| s(Time):NH speakers | 0.50911 | 9 | 0.108 | **7.44e-05** |
| s(Time,Speaker) | 494.11704 | 868 | 4.646 | **<2e-16** |

**Table S2:** Results from individual one sample t tests for Experiment 2, showing differences from 0.

|  |  | early window |  |  |  |  |  | late window |  |  |  |  |
| --- | --- | --- | --- | --- | --- | --- | --- | --- | --- | --- | --- | --- |
|  |  | *M* | *SD* | *df* | *t* | *p* | *d* | *M* | *SD* | *t* | *p* | *d* |
| up | NH | 14.76 | 15.91 | 46 | 6.37 | **<.001** | 0.93 | 35.90 | 25.00 | 9.84 | **<.001** | 1.44 |
|  | CA | 11.42 | 16.31 | 40 | 4.48 | **<.001** | 0.70 | 21.05 | 25.33 | 5.32 | **<.001** | 0.83 |
| down | NH | 19.46 | 17.26 | 46 | 7.73 | **<.001** | 1.13 | 42.18 | 29.43 | 9.83 | **<.001** | 1.43 |
|  | CA | 13.99 | 20.49 | 40 | 4.37 | **<.001** | 0.68 | 30.22 | 30.61 | 6.32 | **<.001** | 0.99 |

**Table S3:**
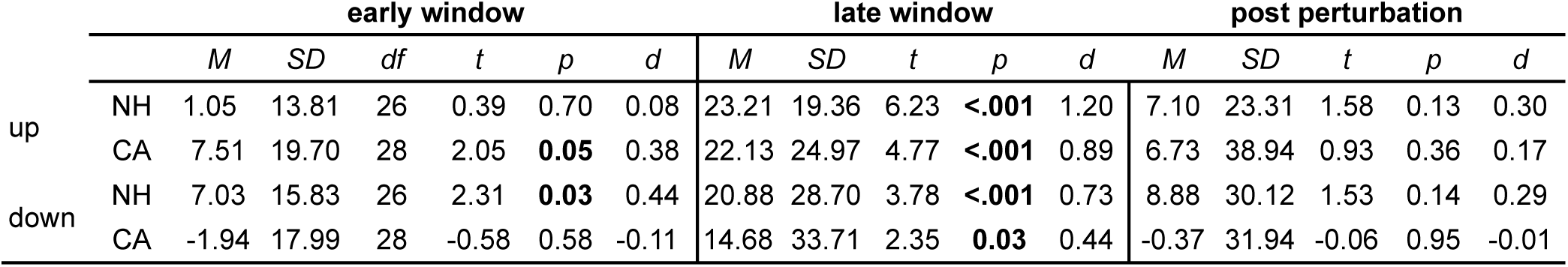
Results from individual one sample t tests for Experiment 3, showing differences from 0.

|  |  | early window |  |  |  |  |  | late window |  |  |  |  | post perturbation |  |  |  |  |
| --- | --- | --- | --- | --- | --- | --- | --- | --- | --- | --- | --- | --- | --- | --- | --- | --- | --- |
|  |  | *M* | *SD* | *df* | *t* | *p* | *d* | *M* | *SD* | *t* | *p* | *d* | *M* | *SD* | *t* | *p* | *d* |
| up | NH | 1.05 | 13.81 | 26 | 0.39 | 0.70 | 0.08 | 23.21 | 19.36 | 6.23 | **<.001** | 1.20 | 7.10 | 23.31 | 1.58 | 0.13 | 0.30 |
|  | CA | 7.51 | 19.70 | 28 | 2.05 | **0.05** | 0.38 | 22.13 | 24.97 | 4.77 | **<.001** | 0.89 | 6.73 | 38.94 | 0.93 | 0.36 | 0.17 |
| down | NH | 7.03 | 15.83 | 26 | 2.31 | **0.03** | 0.44 | 20.88 | 28.70 | 3.78 | **<.001** | 0.73 | 8.88 | 30.12 | 1.53 | 0.14 | 0.29 |
|  | CA | -1.94 | 17.99 | 28 | -0.58 | 0.58 | -0.11 | 14.68 | 33.71 | 2.35 | **0.03** | 0.44 | -0.37 | 31.94 | -0.06 | 0.95 | -0.01 |

**Table V:** Results from individual one sample t tests for Experiment 4, showing differences from 0.

|  |  | early window |  |  |  |  |  | late window |  |  |  |  |
| --- | --- | --- | --- | --- | --- | --- | --- | --- | --- | --- | --- | --- |
|  |  | *M* | *SD* | *df* | *t* | *p* | *d* | *M* | *SD* | *t* | *p* | *d* |
| up | NH | 13.19 | 12.24 | 16 | 4.44 | **<.001** | 1.08 | 29.50 | 18.92 | 6.43 | **<.001** | 1.56 |
|  | CA | 14.27 | 18.73 | 14 | 2.95 | **0.01** | 0.76 | 21.90 | 30.00 | 2.83 | **0.01** | 0.73 |
| down | NH | 15.90 | 14.32 | 16 | 4.58 | **<.001** | 1.11 | 25.81 | 18.76 | 5.67 | **<.001** | 1.38 |
|  | CA | 13.05 | 21.72 | 14 | 2.33 | **0.04** | 0.60 | 15.49 | 28.77 | 2.09 | 0.06 | 0.54 |

